# Morphological aspects of immature stages of *Migonemyia migonei* (Diptera: Psychodidae, Phlebotominae) an important vector of Leishmaniosis in South America by scanning electron microscopy

**DOI:** 10.1101/2020.07.22.215624

**Authors:** Eric F. Marialva, Nágila F. Secundino, Fernando F. Fernandes, Helena R. C. Araújo, Claudia M. Ríos-Velásquez, Paulo F. P. Pimenta, Felipe A. C. Pessoa

**Affiliations:** Laboratório de Ecologia de Doenças Transmissíveis na Amazônia, Instituto Leônidas & Maria Deane – Fiocruz Amazônia, Manaus, Amazonas, Brazil; Laboratório de Entomologia Médica, Instituto René Rachou – Fiocruz Minas Gerais, Belo Horizonte, Minas Gerais, Brazil; Programa de Pós Graduação em Biologia da Interação Patógeno Hospedeiro, Instituto Leônidas & Maria Deane – Fiocruz Amazônia, Manaus, Amazonas, Brazil; Division of Entomology, Federal University of Goiás, Goiânia, Goiás, Brazil; Instituto de Ciências Biomédicas, Departamento de Parasitologia, Universidade de São Paulo, São Paulo, Brazil

## Abstract

We describe the immature stages of *Migonemyia migonei*, which is the vector of *Leishmania (Viannia) braziliensis*, the aetiological agent of cutaneous leishmaniasis in South America, and a putative vector of Leishmania infantum chagasi. Scanning Electron Microscopy (SEM) was used to refine the description of the structures of eggs, all instar larvae and pupa. The eggs have polygonal cells on the egg exochorion, and differences between larval and pupal chaetotaxy are highlighted. Different sensillary subtypes were observed in the larval stages, among the types trichoidea, basiconica, coelonica and campanoformia. These results contribute to the taxonomy of *Mg. migonei* and may contribute to future studies on the phylogeny of this important species vector.

## Introduction

In the last few decades, new proposals in phylogeny sand flies, especially based on adult morphology has been highlighted and several authors have adopted the Galati [1,2] proposal, that changed the taxonomy, phylogeny and nomenclature basis of phlebotomine systematic. However, the knowledge about several aspects of the immature stages of phlebotomine sand flies (Diptera: Psychodidae) is still a challenge, due to the difficulties of finding a natural breeding site or lab colonization. So, of about more than 537 species described in the Neotropics [1–3], larval stages or mature larvae and rarely pupae of only 92 species of the New World sand flies have been described, or partially described.

Larval structures are important for providing some highlights on taxonomy, phylogeny, and evolution of this subfamily. In the last few decades, new proposals have been highlighted on phylogeny of sand flies, especially based on adult morphology, and several authors have adopted the Galati proposal [1,2], which has changed the taxonomy, phylogeny and nomenclature basis of phlebotomine systematic.

The analysis of the microstructure of the immature stages of Phlebotominae, in addition to contributing to the discovery of other morphological characters capable of promoting taxonomic and phylogenetic studies, in order to better elucidate the evolution of this subfamily, also makes it possible to investigate the existence of sense structures used in the communication of these vectors, aiming at the development of alternative eco-friendly control strategies.

The use of scanning electron microscopy (SEM) has significant improved the characterization and descriptions of immature forms, and provided details of larval chaetotaxy [4,5]; ontogeny [6–8]; spiracles [9,10]; antennal, and mouthparts, such as the sensilla as well as caudal bristles [7,11,12]. Despite this, only a few articles have been carried out about the pupal morphology of New World phlebotomine sand flies [8,13–16]. Therefore, the number of descriptions of immature forms of sand flies still remains scarce.

The sand fly, *Mg. migonei* (França), is an important vector of *Leishmania (Viannia) braziliensis*, and one of the causative agents of cutaneous leishmaniosis in South America, especially in Brazil [17–19]. Torrellas [20] found *Mg. migonei* infected with *Le. guyanensis* and *Le. mexicana* in an Andean region of Venezuela. Studies confirmed that this species is also associated with the transmission of *Leishmania infantum chagasi* in Brazil and Argentina [21,22].

Despite the importance of its immature morphology, few studies have been carried out, especially under scanning electron microscopy (SEM). These were restricted to the description of the larvae antennae [11] and spiracles [9,10] and to the egg exochorion [23,24].

The present study aims to provide a complete morphological analysis of the surface of immature stages of *Mg. migonei*, in order to reveal taxonomic characters that can support future works on phylogenetics and systematics involving immature stages of this vector.

## Material and Methods

The eggs, larvae and pupae of *Mg. migonei* were acquired from a stable colony maintained in laboratory conditions, whose parents were obtained in the municipality of Baturité, Ceará state, Brazil. The species was bred at the laboratory facilities of Leônidas & Maria Deane Institute, Manaus, Amazonas state, according to the method described by Lawyer [25]. Some of the larvae from each larval instar (1^st^ to 4^th^) and pupae were slide-mounted in Berlese fluid. Measurements of the body’s bristles were made under eye pierce using light microscopy.

Morphology and chaetotaxy of the head were observed following the methodology of Arrivillaga[26], which indicates the morphology and setae of the mouthparts with taxonomical importance. Chaetotaxy of the body followed the system used by Ward [27]. The chaetotaxy of the pupae used in this study followed the terminology proposed by Oca-Aguilar [8]. Systematic classification follows that proposed by Galati [2], and abbreviations of the genera follow Marcondes [28]. In addition, both species were studied and photographed under scanning electron microscopy. Some reared larvae were killed in hot water (70°C), fixed in 3% glutaraldehyde and then washed thoroughly in phosphate-buffered saline; and the solution was changed every 30 min over a period of six hours. Subsequently, they were fixed in osmium tetroxide, dehydrated in a series of ethyl alcohol concentrations, submitted to critical point drying in carbon dioxide and spattered with 25 MA colloidal gold [7,11]. The specimens were examined in a scanning electron microscope (JSM5600, JEOL, Tokyo, Japan) at an accelerating voltage of 7 KV and then photographed. Tables were mounted showing the differences in chaetotaxy between the instars of each species and between species.

## Results

### Egg of *Mg. migonei*

The egg is elongated, with one side slightly flattened, measuring 323 (300-351) μm in length and 94.8 (89-107) μm in width (N=4) (Fig.1A). The exochorion is formed by a thin basal lamina that supports their ornaments or sculptures with polygonal reticulation, which comprises ridges, usually continuous, forming alternating transversal rows of generally rectangular parallel cells or square to polygonal cells (Fig. 1B).

**Fig. 1.**
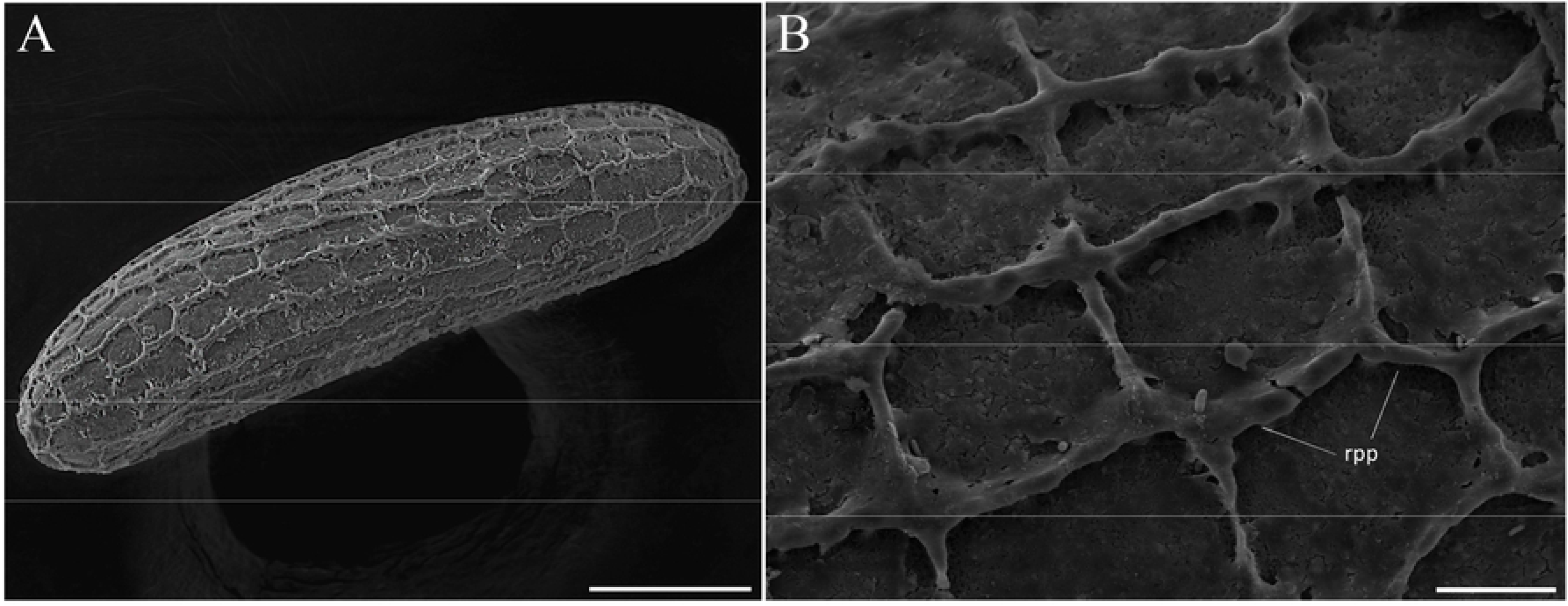
Scanning electron microscopy of the eggs of *Migonemyia migonei*. A, general view of the egg showing an ornamentation characterized by the presence of ridges arranged in a polygonal pattern (rpp; scale bar: 50 μm); B, eggshell ornamentation showing detail of the ridges (Scale bar: 10 μm).

### General appearance of the larvae of *Mg. migonei*

The larva is caterpillar-like, with a well-sclerotized hypognathous, non-retractile head with very short antennae with short basal tubercle. The dark brownish colour head and body tegument are covered by very small spines and tubercles in a scattered distribution. The thorax includes prothorax, with the anterior spiracle borne laterally, and other two segments (meso and metathorax). Its abdomen is nine-segmented, covered by brown pale setae and the body tegument is yellowish, with a pair of posterior spiracles borne laterally on a short tubercle.

Caudal filaments (or caudal setae), long sensilla of the trichoid type that exhibit many wall pores, implanted between non-parallel ridges and that interconnect (or which overlap), darkened, were observed double-paired in the last three instars, or simply paired when is in the first instar (Fig. 2A-B).

**Fig 2.**
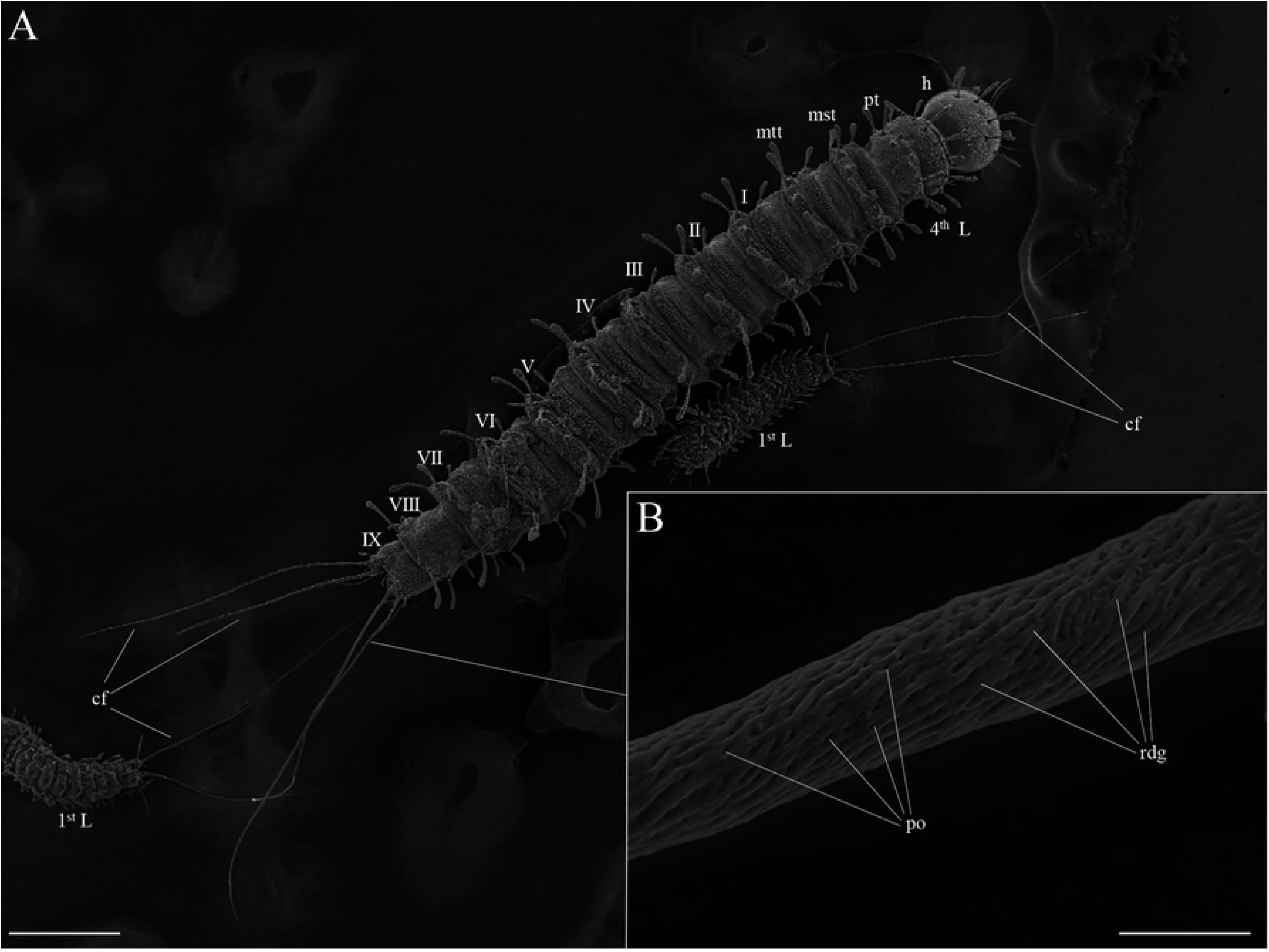
Scanning electron microscopy of the fourth, and first instar larva of *Migonemyia migonei*. A, observation of first (1st) and fourth (4th) instar larvae, besides of the number of body segments and caudal filaments (scale bar: 200μm); B, higher magnification of the surface of the caudal filament showing pores implanted between non-parallel, interconnected and overlapping ridges. (scale bar: 5μm). Caudal filaments (cf), head (h), protorax (pt), mesotorax (mst), metatorax (mtt), ridges (rdg), and pores (po).

The head is dark brown (Fig. 2A, 3A), body colour is pale with darkened eighth and ninth abdominal segments and bears tiny spines in all segments (Fig. 2A, Fig 7), the first instar is present a prominent egg buster (Fig 3C-D) with peculiar shape. There are three types of setae, usually distributed in pairs: a barbed brush-like setae (of brush-like trichoid sensilla type), more widely distributed on the larval head and body (Fig.3A, seta 2) and, a little barbed (weakly brush-like trichoid sensilla; Fig. 3A, seta 1) and a simple, bare paired setae (trichoid sensilla; Fig. 3A, seta 6). The size and the type of setae are shown in Table 1.

**Fig 3.**
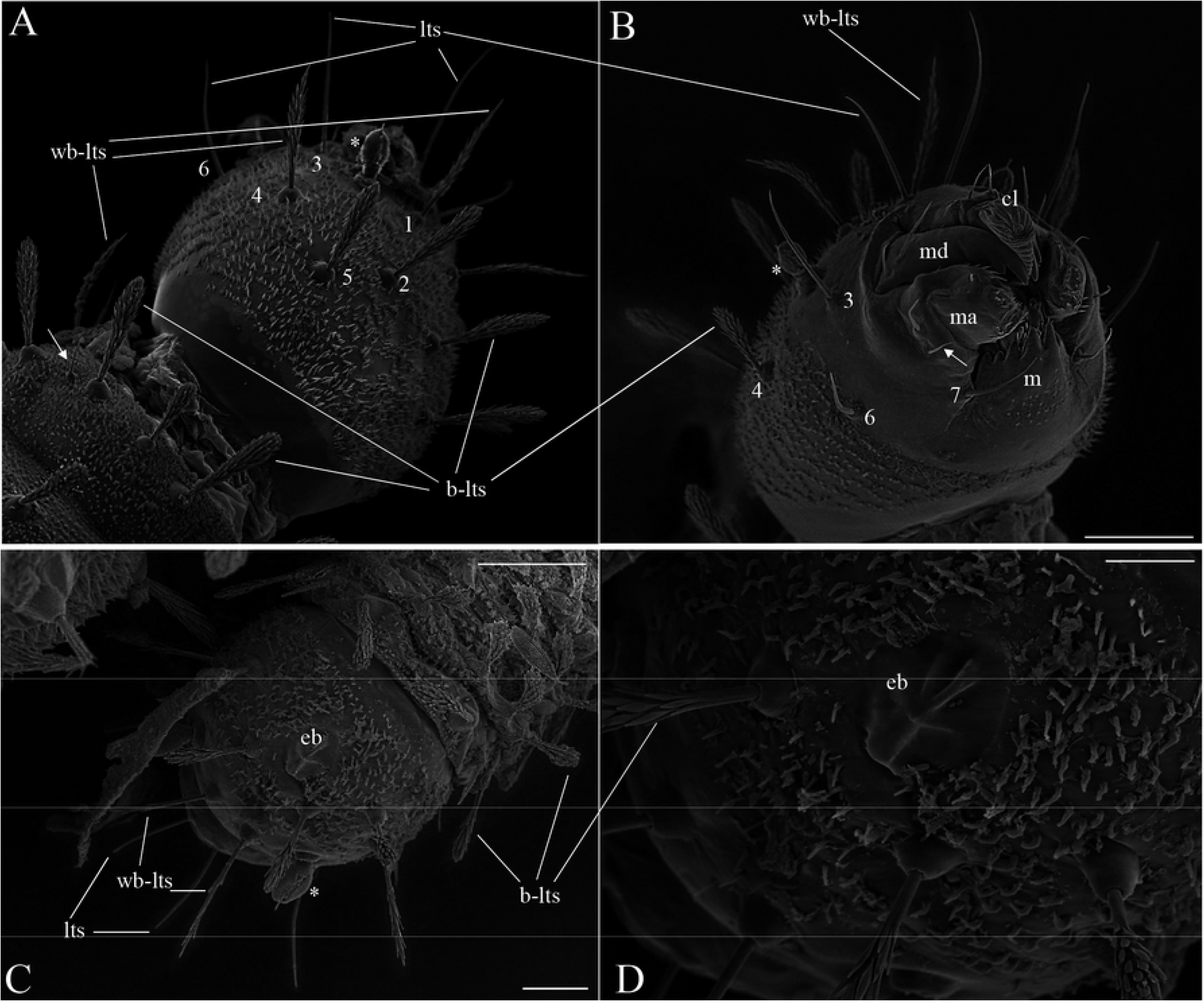
Scanning electron microscopy of the larva of *Migonemyia migonei*. A, Head of the fourth instar larval in dorsal and ventral view B. Long trichoid sensilla observed at the apex of the head (lts) and the short trichoid sensilla on the mouthparts (arrows; A and B scale bars: 20 μm); C, Head in dorsal of the first. On the forehead are two weakly brush-like trichoid sensilla (wb-lts) inserted slightly forward and between the antennae (ant) and long trichoid sensilla (lts) are inserted furtherdown toward the mouthparts (scale bar: 20 μm). D, Egg buster (eb). cl, clipeo; md, mandible; ma, maxilla; m, mentum. The setae were numbered according to the chaetotaxy proposed in study: (1 = wb-lts) frontoclipeal anterior setae (weakly brush-like trichoid sensilla); (2 = b-lts) frontoclipeal posterior setae with barbed shape (brush-like trichoid sensilla); (3 = lts) the genal anterior setae with a simple spine form (long and bare trichoid sensilla); the genal medial (4) and genal posterior (5) are barbed brush-like setae (= brush-like trichoid sensilla). In the ventral part (Fig. 3B), the postgenal (6) and subgenal (7) are simple setae (bare trichoid sensilla)

**Table 1.** Corresponding numbers and size of the setae of each segment (size in μm, N=4) of fourth to first instar larvae of *Migonemyia migonei*.

| Larva body part | *Migonemyia migonei* |  |  |  |  |  |
| --- | --- | --- | --- | --- | --- | --- |
|  | Setae number | Type of setae | Size of the setae in the corresponding larval instar |  |  |  |
| Head |  | Bristle | 4<sup>th</sup> | 3<sup>rd</sup> | 2<sup>nd</sup> | 1<sup>st</sup> |
| Frontoclipeal anterior | 1 | Spinulate | 93.3 | 77.4 | 65.4 | 52.0 |
| Frontoclipeal posterior | 2 | Barbed | 85.3 | 65.3 | 50.7 | 29.4 |
| Genal anterior | 3 | Simple | 113.0 | 70.65 | 60.0 | 41.4 |
| Genal medial | 4 | Barbed | 93.3 | 64 | 53.3 | 44.0 |
| Genal posterior | 5 | Barbed | 90.7 | 62.7 | 46.7 | 33.4 |
| Postgenal | 6 | Simple | 101.3 | 64.0 | 54.7 | 37.3 |
| Subgenal | 7 | Simple | 80.0 | 37.3 | 24.0 | 13.3 |
| Prothorax |  |  |  |  |  |  |
| Dorsal internal | 1 | Barbed | 85.0 | 57.3 | 46.7 | 38.7 |
| Dorsal intermediate | 2 | Barbed | 65.0 | 41.35 | 20.0 | 24.0 |
| Dorsal external | 3 | Barbed | 87.5 | 60.0 | 49.4 | NO |
| "Shoulder" accessory | a | Spine | 25.0 | 17.4 | 17.7 | NO |
| "Shoulder" accessory | b | Barbed | 16.0 | 08.0 | 08.0 | NO |
| Anterior ventrolateral | 4 | Barbed | 100.0 | 61.3 | 45.4 | 32.0 |
| Ventral external | 5 | Barbed | 67.5 | 55.0 | 49.4 | 30.7 |
| Ventral internal | 6 | Barbed | 100.0 | 64 | 53.3 | 29.3 |
| Dorsal submedian | 7 | Barbed | 90.0 | 49.3 | 33.4 | 16.0 |
| Mid-dorsal | 8 | Barbed | 100.0 | 64.0 | 44.0 | 18.7 |
| Dorsolateral | 9 | Barbed | 95.0 | 52.0 | 40.0 | 26.7 |
| Basal | 10 | Barbed | 42.5 | 22.7 | 12.0 | NO |
| Post-ventrolateral | 11 | Barbed | 96.0 | 53.35 | 44.0 | 30.7 |
| Post-ventral | 12 | Spine | 13.3 | 6.7 | 6.7 | 6.7 |
| Mid ventral | 13 | Barbed | 72.0 | 42.6 | 26.7 | 12.0 |
| Ventral intermediate | 14 | Barbed | 17.3 | 12.0 | NO | 2.7 |
| Ventral submedian | 15 | Barbed | 38.7 | 21.35 | 13.3 | 8.0 |
| Meso and metathorax |  |  |  |  |  |  |
| "Shoulder" accessory | a | Spine | 14.7 | 10.7 | 8.0 | 6.7 |
| "Shoulder" accessory | b | Barbed | 17,3 | 9,35 | 8.0 | NO |
| Anterior ventrolateral | 4 | Barbed | 90.0 | 52.0 | 37.4 | 25.4 |
| Dorsal submedian | 7 | Barbed | 142.5 | 69.35 | 37.4 | 20.0 |
| Mid-dorsal | 8 | Barbed | 152.5 | 78.65 | 46.7 | 21.4 |
| Dorsolateral | 9 | Barbed | 127.5 | 65.3 | 42.7 | 29.3 |
| Basal | 10 | Barbed | 25.0 | 13.3 | 8.0 | NO |
| Post-ventrolateral | 11 | Barbed | 78.8 | 49.3 | 36.0 | 17.4 |
| Post-ventral | 12 | Spine | 14.7 | 9.35 | NO | 05.3 |
| Mid-ventral | 13 | Barbed | 77.3 | 38.65 | 29.3 | 14.7 |
| Ventral intermediate | 14 | Barbed | 22.7 | 13.3 | 8.0 | 4.0 |
| Ventral submedian | 15 | Barbed | 44.0 | 21.4 | 14.7 | 9.4 |
| Abdominal segments 1–7 |  |  |  |  |  |  |
| Dorsal intermediate | 2 | Barbed | 35.0 | 12.0 | 5.33 | 2.7 |
| Anterior ventrolateral | 4 | Barbed | 102.5 | 48.0 | 34.65 | 16.0 |
| Dorsal submedian | 7 | Barbed | 165.0 | 76.0 | 38.70 | 17.4 |
| Mid-dorsal | 8 | Barbed | 180.0 | 93.3 | 50.65 | 20.0 |
| Dorsolateral | 9 | Barbed | 165.0 | 82.65 | 48.0 | 41.4 |
| Post-ventrolateral | 11 | Barbed | 75.0 | 44.0 | 36.0 | 18.7 |
| Post-ventral | 12 | Barbed | 30.0 | 13.3 | 10.7 | NO |
| Ventral submedian | 15 | Simple | 66.7 | 37.3 | 32.0 | 17.4 |
| “C”- |  | Spine | 12.0 | 9.35 | 8.00 | NO |
| Abdominal segment 8 |  |  |  |  |  |  |
| Anterior ventrolateral | 4 | Barbed | 77.3 | 34.7 | 25.4 | NO |
| Dorsal submedian | 7 | Barbed | 46.7 | 20.0 | 10.7 | 10.7 |
| Mid-dorsal | 8 | Barbed | 147.15 | 88.0 | 57.3 | 12.0 |
| Dorsolateral | 9 | Barbed | 118.8 | 69.4 | 45.3 | 37.3 |
| Post-ventrolateral | 11 | Spinulate | 52.0 | 28.0 | 25.4 | 12.0 |
| Post-ventral | 12 | Spine | 22.0 | 12.0 | 6.7 | 8.0 |
| Ventral submedian | 15 | Spinulate | 57.3 | 26.7 | 25.4 | 9.35 |
| “Should accessory” | A | Spine | 14.7 | 6.7 | 5.4 | NO |
| “Should accessory” | B | Spine | 29.3 | 12.0 | 9.4 | 5.3 |
| Abdominal segment 9 |  |  |  |  |  |  |
| Anterior ventrolateral | 4 | Simple | 220 | 152 | 137.5 | 92.0 |
| Dorsal submedian | 7 | Simple | 95.9 | 64.0 | 46.7 | 34.7 |
| Mid-dorsal | 8 | Barbed | 57.3 | 33.4 | 24.0 | 52.0 |
| Dorsolateral | 9 | Barbed | 57.3 | 36.0 | 24.0 | 16.0 |
| Post-ventrolateral | 11 | Simple | 97.2 | 48.0 | 38.7 | 21.3 |
| Post-ventral | 12 | Simple | 48.0 | 22.7 | 21.3 | 9.35 |
| Ventral submedian | 15 | Simple | 37.3 | 21.3 | 14.7 | 13.3 |
| Internal caudal | IC | Simple | 1250 | 950 | 720 | 560 |
| External caudal | EC | Simple | 1045 | 625 | 565 | NO |

NO: Not observed

### Head

The head is capsule-like, broader than it is high. The tegument is covered by thin, small spicules of the microthrichia type. On the dorsal part of the head (Fig.3A), the cephalic tagma has the following setae: the anterior frontoclypeal setae (1 weakly brush-like trichoid sensilla, subtype) with spinulate form, the posterior frontoclypeal setae (2;brush-like trichoid sensilla, subtype) with barbed shape, the anterior genal setae (3) with a simple spine form. The medial genal (4) and posterior genal (5) setae are barbed brush-like (brush-like trichoid sensilla, subtype). In the ventral part (Fig. 3B), the postgenal (6) and subgenal (7) are simple setae (Table 1). All setae are inserted in small tubercles. In the first instar larvae, the setae 1 is usually simple; however, it is possible to predict the projection of those setae becoming barbed at the other subsequent instars. Each of the antennae of *Mg. migonei* larvae (Fig. 4A-B) has a basal tubercle (socket) that is a small and cylindrical segment fused at a second ovoid distal segment. This segment presents an antennal organ, which is equipped with a longitudinal furrow in the posterior surface, and is more evident in the 1^st^ instar antennae, as well as three short structures in the base of the segment. The central structure is wider than it is long and shorter than the laterals ones (Figs. 4A-B). The apex of antennae exhibits a single apical clavate basiconic sensillum and, from a lateral groove, four sensilla of the *coeloconica* type emerge: three smaller with blunt apex – noting that the middle one (smaller coeloconic sensillum, subtype) is wider, shorter and less cylindrical than the two that are on the sides (blunt coeloconic sensilla, subtype) – and, between these three, behind, one larger and clavate (clavate coeloconic sensillum, subtype; Figs. 4A-B).

**Fig 4.**
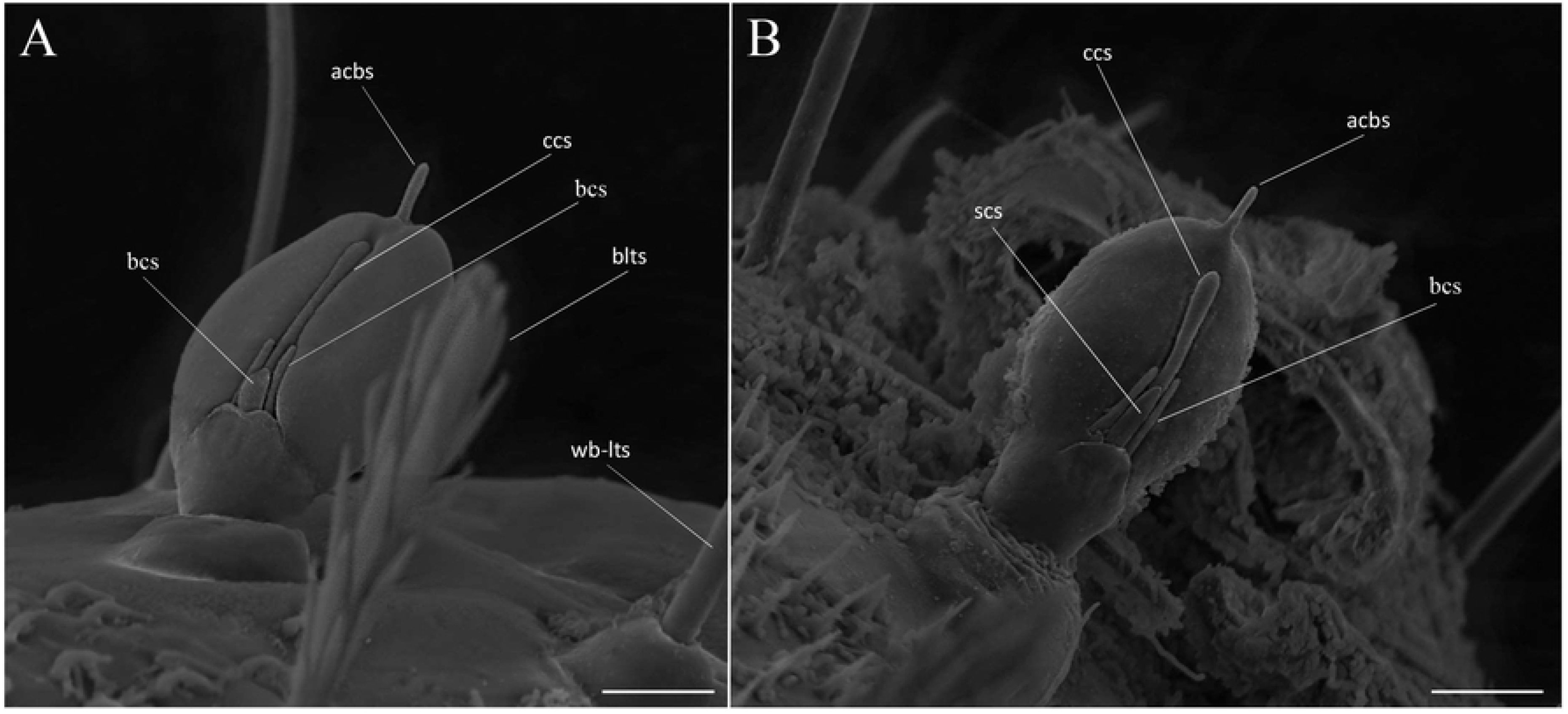
Scanning Electron Microscopy of the larva of *Migonemyia migonei*. A, antenna of the first instar larva (scale bar: 5 μm); B. antenna of the fourth instar larva (scale bar: 10 μm). It is observed in A and B, a single apical clavate basiconic sensillum (acbs) at the apex of the antennae and, emerging from a lateral groove, one smaller coeloconic sensillum (scs) and two blunt coeloconic sensilla (bcs) and, between these two, behind, one clavate coeloconic sensillum (ccs).

Mouthparts: The external part of the mouth is composed of a pair of mandibles, a pair of maxillae, labrum, and mentum. Each segmented mandible bears two simple setae in the middle of the dorsal part (S1 and S2), and a simple seta (S6) in the superior margin of the mandible, which are similar to those described by Pessoa et al. (2008). In the lower part of the mandible, there are five strong teeth, a proximal (T3) nearest the molar lobe (ML), an apical single (T1), and double-paired median tooth (T2 – that are observed, in light microscope, as a single tooth) (Fig. 6).

Each maxilla has three simple setae, an S1 in the apical dorsal part, and two (S2 and S3) in the proximal part (Fig. 5). There is a maxillary process in the middle of this structure. In the margin of the dorsal part, there is a sequence of a small and sparse comb of spines, similar to those found in the maxilla of *Ev. lenti* and *Ev. carmelinoi* [12]. At the apex, there are papilliform and trichodea sensillae (spinous hairs). On the upper side, there is a row of small setae. The ventral surface of the labrum is covered with parallel, transverse rows of finger-like combs of setae; the dorsal side has two pairs of very small simple setae. The clypeus has two pairs of simple setae, the distal pair is small, and the apical bigger (Fig. 5).

**Fig 5.**
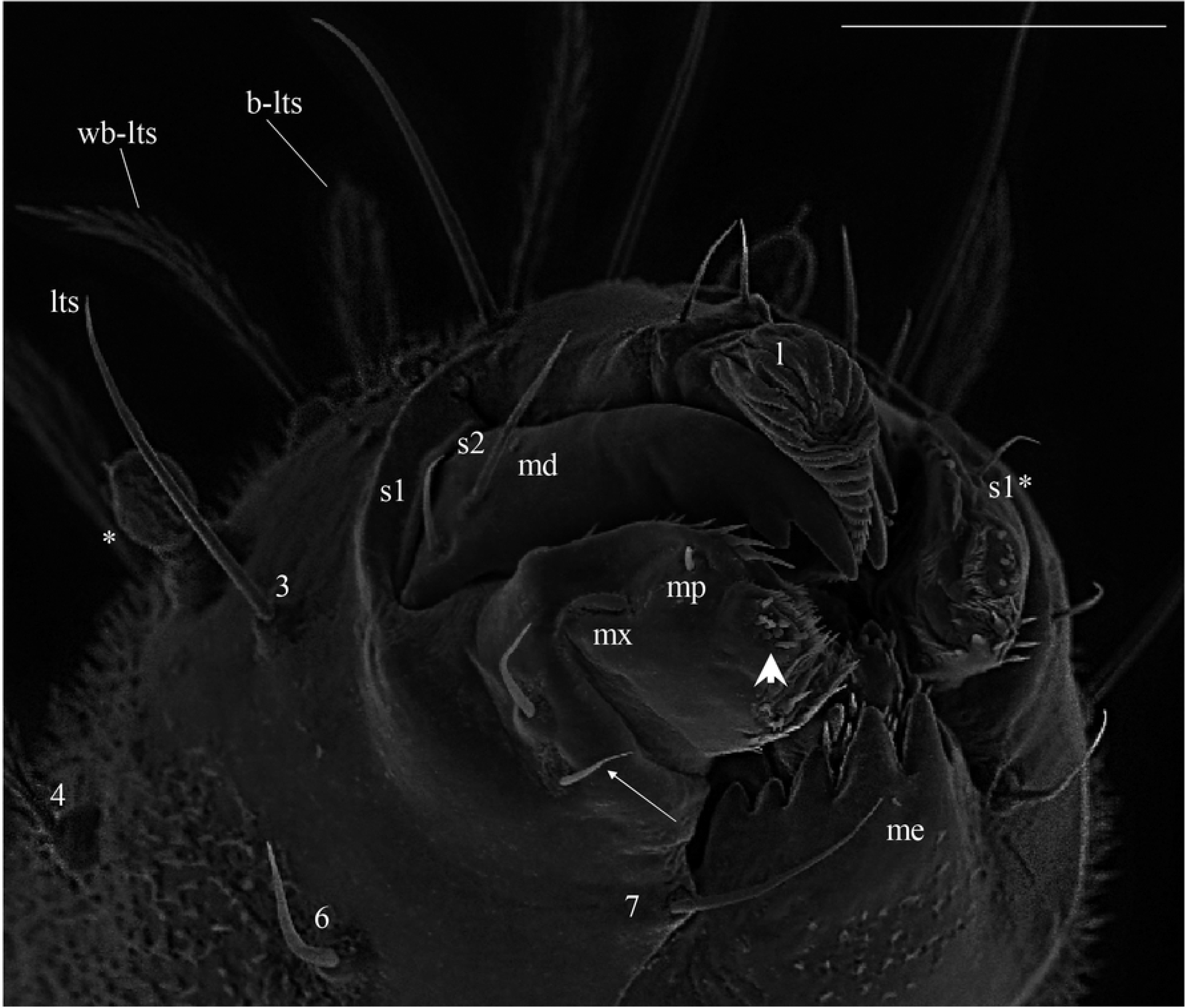
Scanning electron microscopy of the mouthparts of fourth instar larva of *Migonemyia migonei*. L, labrum; md, mandible; mx, maxila; me, mentum; s1-s6, mandible setae; s1*-s3*, mx : maxila setae; maxillary palpus (scale bar: 20 μm).

**Fig 6.**
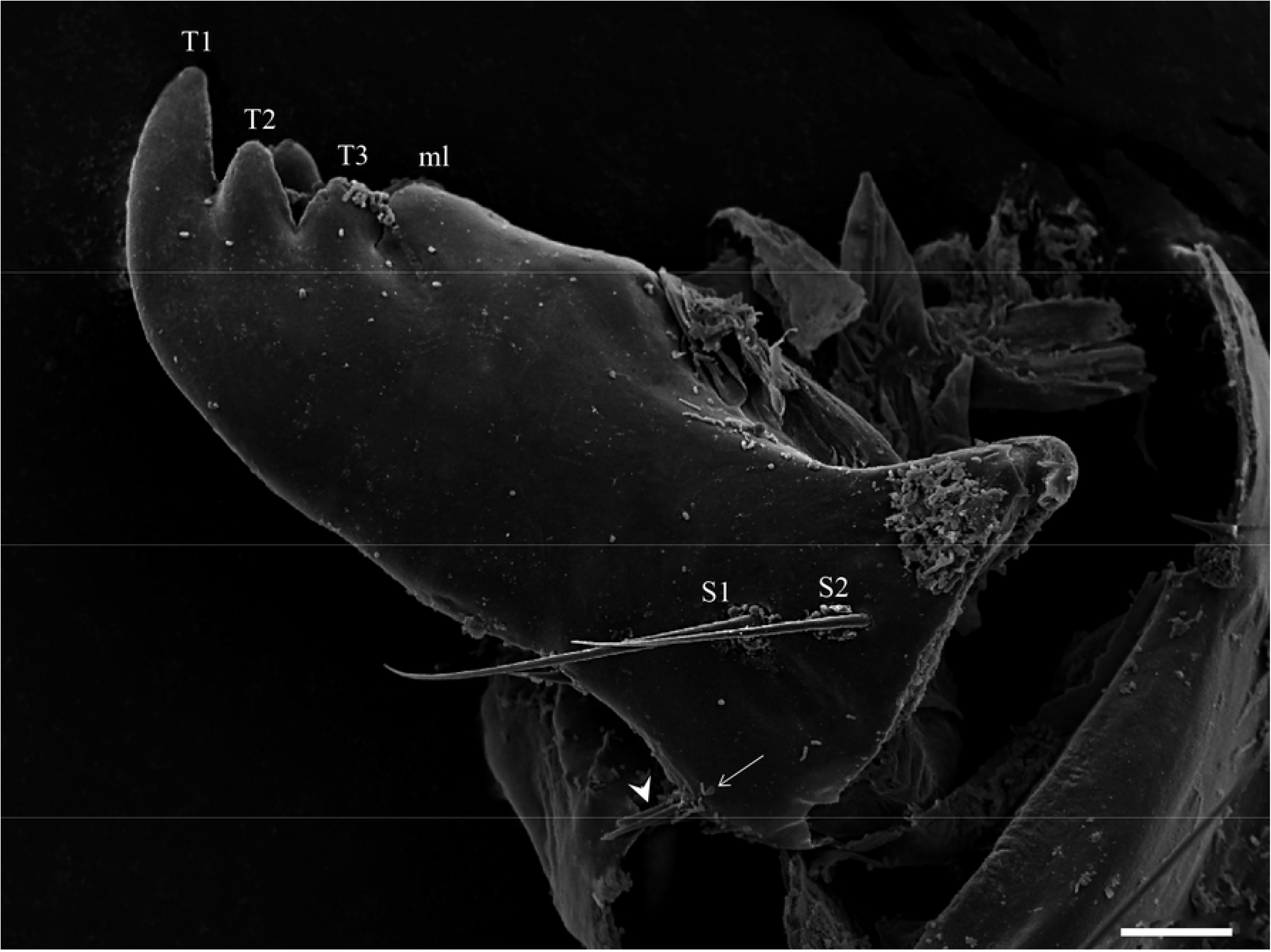
Mandible of *Migonemyia. migonei* larva. Apical (T1), double paired median tooth (T2), proximal (T3) and molar lobe (ml). Proximally, setae of the trichoid sensilla type (S1 and S2) are observed and, below these, a small seta (arrow) and, under the margin, the ends of three other setae (arrowhead; scale bar:10 μm).

The thorax has three segments, the prothorax has the appearance of two segments, and the meso and metathorax are homologous with the posterior setae of the prothorax. Chaetotaxy follows the same pattern of setae as classified by Ward [27] and is presented in Table 1, and Figures 7 and 8. The anterior spiracles are conical and have eight to nine papillae (Fig. 9A), though five to six in the third instar larvae, and 4 in the second instar larva (Fig. 9B). However, we did not obtain clear images of the first instar larva in order to compare or count.

**Fig7.**
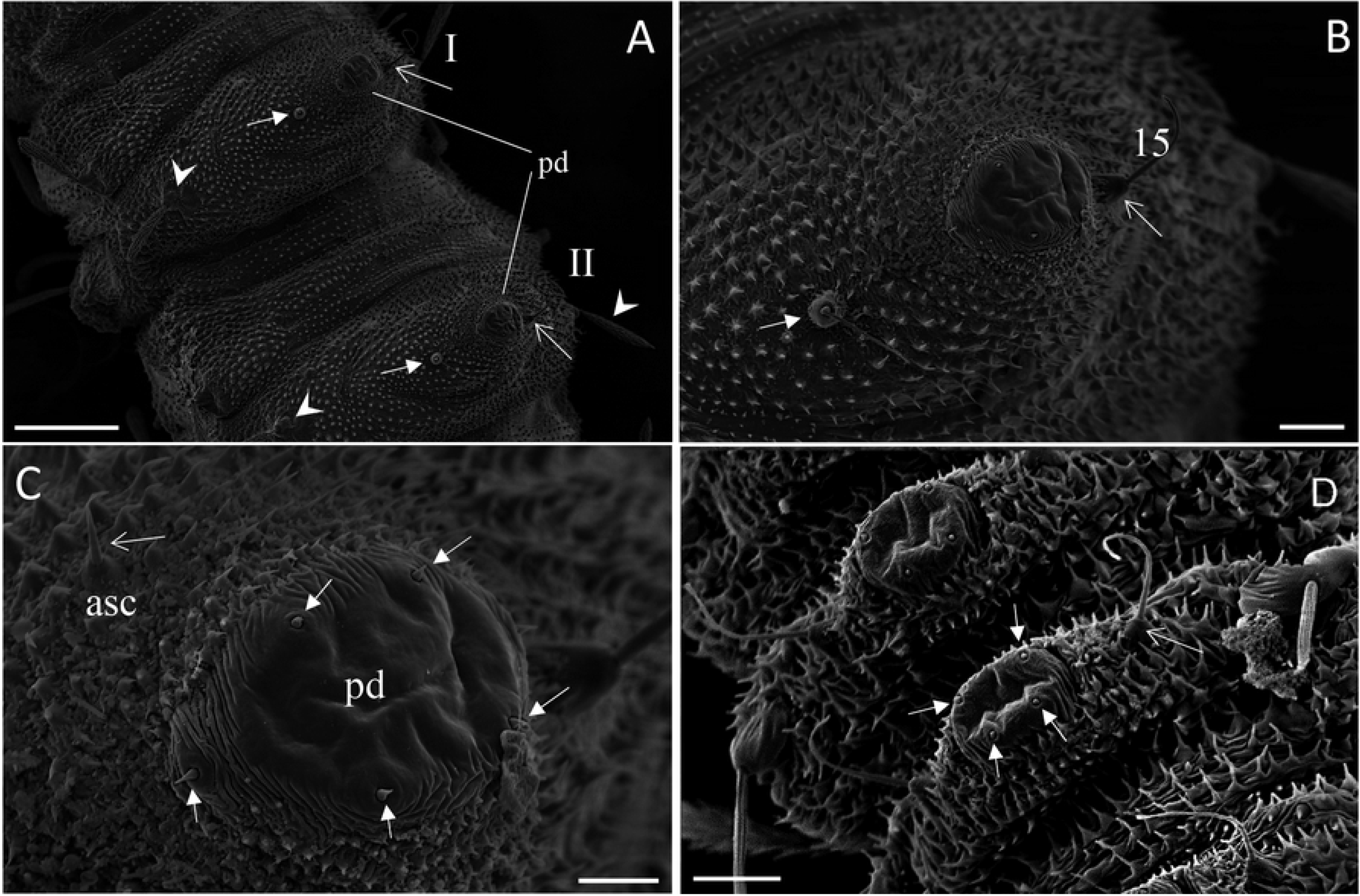
Scanning electron microscopy of the ventral part of abdomen and pseudopoda of *Migonemyia migonei*. A-C, mature larvae; D, first instar larvae. asc: accessory setae, pd: pseudopod, * ampliation accessory setae. (scale bars: 100 μm, 20 μm, 10 μm and 10 μm, respectively).

**Fig 8.**
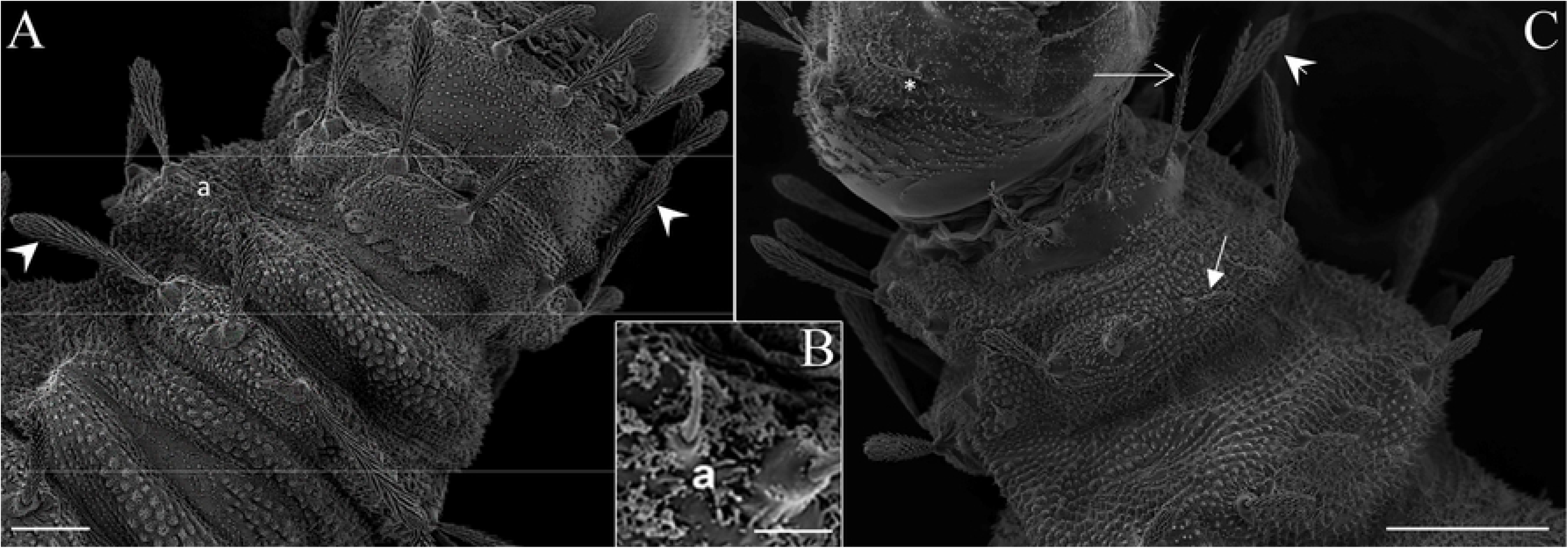
Scanning Electron Microscopy of the mature larva of *Migonemyia migonei*. A, Prothorax and Mesothorax in dorsal of the fourth instar larva; B, a – accessory setae C, – dorsal and ventral view. (scale bars: 50 and 100 μm, respectively).

**Fig 9.**
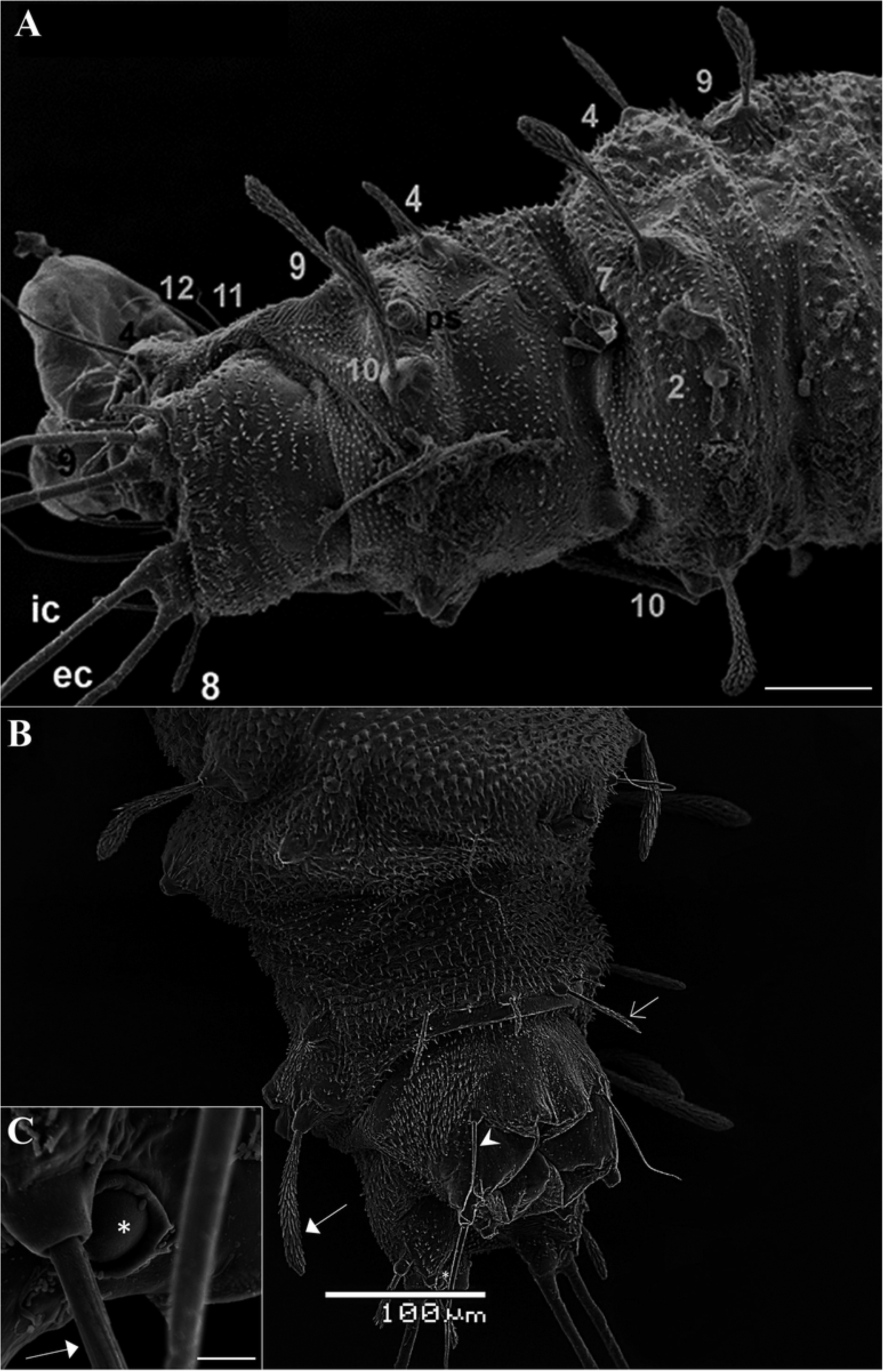
Scanning electron microscopy of the fourth instar larva of *Migonemyia migonei*. A, abdominal 7-9 and in dorsal of the fourth instar; B, and ventral view. Setae numbered according to the chaetotaxy proposed in study; ps – posterior spiracle, *increased area of a campaniform sensilla. Intermediate dorsal (2), Anterior ventrolateral (4), Submedian dorsal (7), Mid–dorsal (8), Dorsolateral (9), Post-ventrolateral (11), Post-ventral (12), Submedian ventral (15). Scale bars: 50 μm and 100 μm, respectively.

**Fig. 10.**
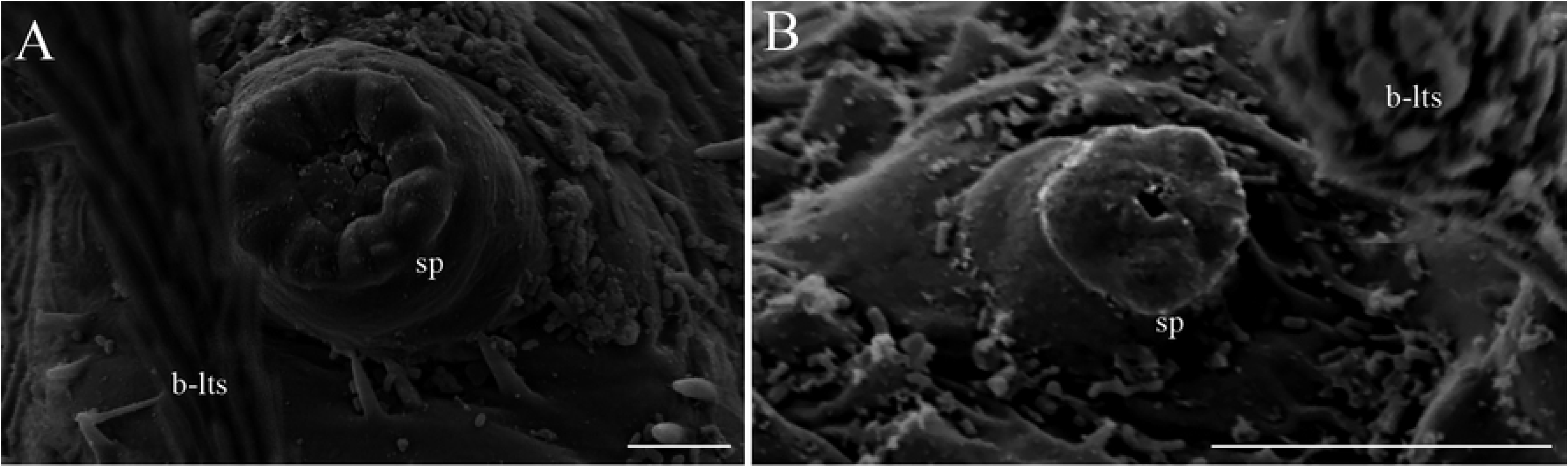
Scanning electron microscopy of the larvae of *Migonemyia. migonei*. A, anterior spiracles of the third instar larva (scale bar: 5 μm); B, anterior spiracles (sp) of the second instar larva brush-like trichoid sensilla.

### Chaetotaxy of prothorax

The tergite has two rows of setae. The first row has three pairs of setae: the dorsal internal, dorsal intermediate and dorsal external, and the second row has two setae, which are the dorsal submedian and the mid-dorsal. The pleura has two setae, the anterior ventrolateral and the dorsolateral, which appear to change position in the larvae. These setae are similar, barbed or brushed-like and have only small differences in size (Table 1). There is a spine hyaline seta, between the first and second rows of setae, usually near the ventrolateral setae. The sternite also has two rows of setae. The first with two similar pairs of setae, a little less barbed than the dorsal setae, the ventral external and the ventral internal. The second row has seven pairs of setae, including the seta b, with different size and shape (Table 1). They are the basal, post-ventrolateral, post-ventral, mid ventral, ventral intermediate, ventral submedian, and b setae. The meso and metathorax do not have the first row of setae that those in the prothorax tergite and sternite have, and the setae are the same as of the second row of setae in the prothorax.

The setae of the abdomen have the same distribution that is proposed by Ward[27], with an absence of the setae number ten in each segment. The segments one to seven are homologous, with similar size and shape. In the anterior part of the pseudopodium, there is a simple pair of seta, which have a similar size to setae eleven and twelve, and these are not considered by other authors as a seta without taxonomic value. This seta is called “c” here (Table 1 and Fig.8C). The eighth and ninth segments are darker. The posterior spiracles are conical, with ten papillae. The shape and measurements of the setae are in Fig. 8A-B and Table 1. Chaetotaxy of the abdominal segments one to seven is as follows: In the tergite, the pairs of setae are not grouped, anterior dorsal intermediate is much smaller than the others, and the anterior ventrolateral is in the border with the pleura, and a row of pairs of setae on the dorsal submedian, mid-dorsal and dorsolateral, also in the border of the pleura. All of them are barbed and have different sizes (Table 1). The sternites (Fig. 8) have large pseudopodia, with a few simple setae, the post-ventrolateral and the post-ventral are both very small, simple setae and a simple and large ventral submedian setae. In the anterior part of the pseudopodia, there is a pair of setae, which is very similar to the post-ventrolateral and the post-ventral, named the setae “c”. The abdominal segments eight and nine lack pseudopodia. Abdominal segment nine ends in two tubercles, each of which bears a caudal filament (Fig. 8A). These tubercles, in the ventral side, also possess a large campaniform sensilla (Fig. 9B). The posterior spiracle has 14-11 papillae.

### Other larval instars

The body sizes of the third to the first instar larvae are (from the head to the end of the ninth abdominal segment and with a maximum width at the metathorax) respectively: 1.7 and 0.27; 0.94 and 0.13; 0.56 and 0.09 mm. The first instar is easily identified by the presence of a unique pair of caudal setae (multiporous long trichoid sensilla; Fig. 2B) and the absence of some bristles in the prothorax (seta 3, a, b and 10) and pro, meso and metathorax (b and 10) and the presence of the egg buster in the head (Fig. 3D). The setae 6 and 14 of the prothorax are simple in this instar and barbed in the others. The setae 11 and 15 of the abdominal segment 8 are simple and, in the other instars, become almost barbed. Chaetotaxy for the other instars are the same of the fourth, but differ in size (Table 1). The pupa emerges from a Y shaped suture of the head of mature larva (Fig 11A-B).

**Fig. 11.**
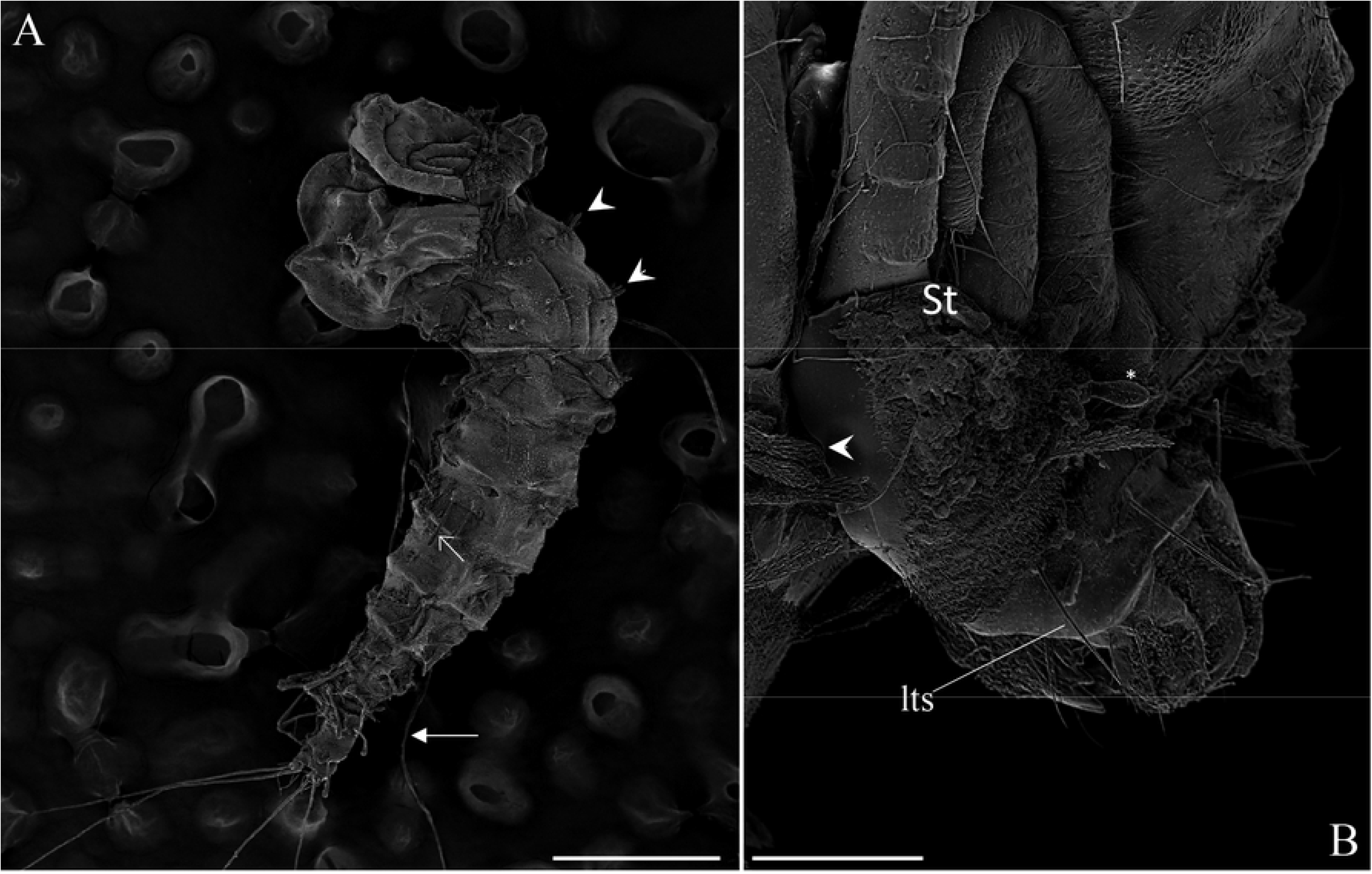
Scanning electron microscopy pupae of *Migonemyia migonei*. A-B emerging from a mature larva; st-suture opened from larva tagma head (scale bars of A and B: 500 and 100 μm, respectively).

### Pupa description

The pupa of *Mg. migonei* is claviform and divided by the cephalothorax and abdomen (Fig. 12A, 13, 15, 16). The pupa tegument has some small ornaments, such as small spines and setae, which are described in Table 3 and the body is covered by several small rounded tubercles. The pharate female pupa is longer (2.24 mm, n = 5) than the pharate males pupa (2.05 mm, n = 5) (Fig. 17).

**Fig 12.**
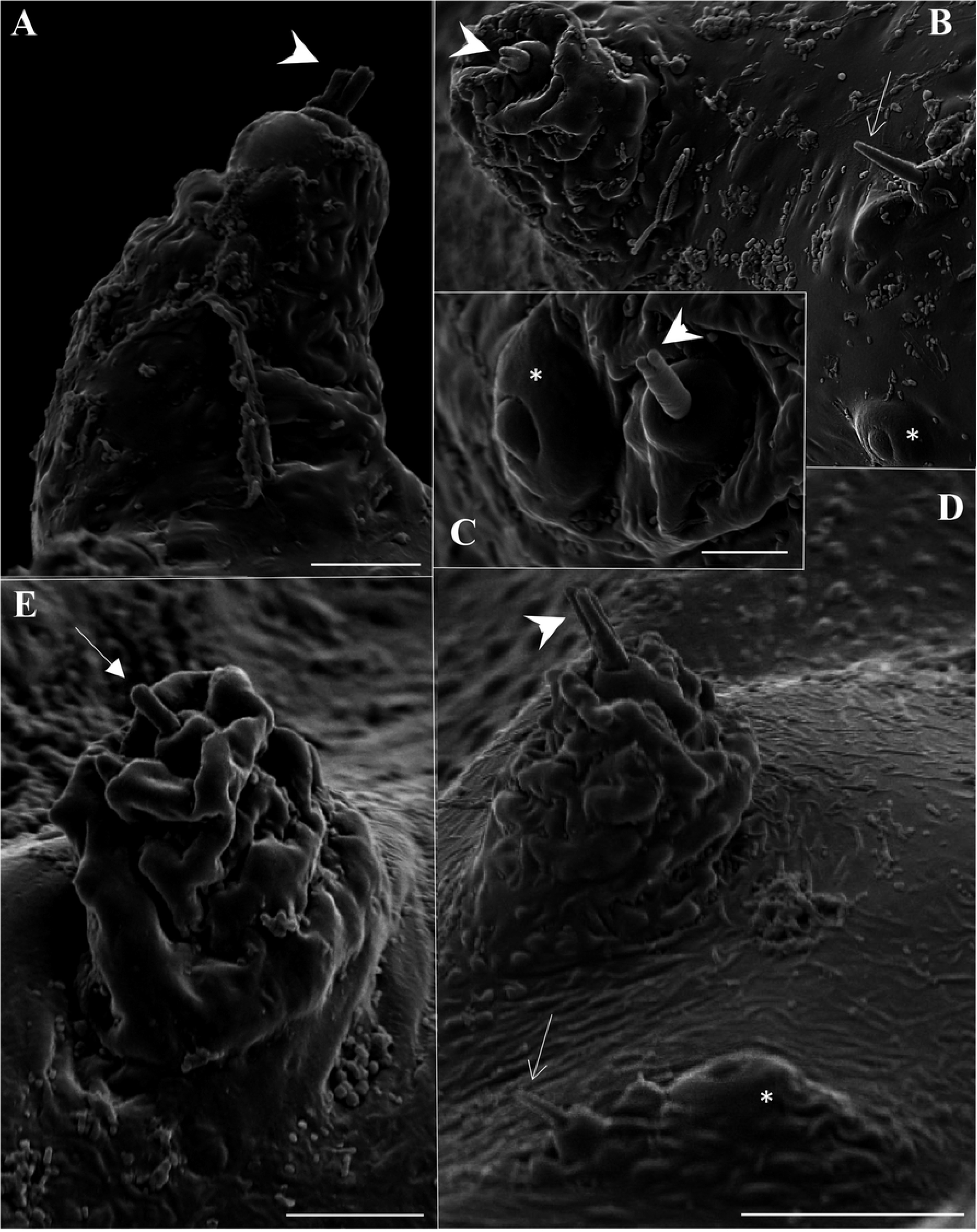
Scanning electron microscopy of the pupa of *Migonemyia migonei*. A, Metathorax setae 1T (scale bar: 10 μm); B, Proothorax setae 1P and 2P (scale bar: 20 μm); C, Metathorax setae 3T (scale bar: 5 μm); D, mesothorax setae 1M and 2M (scale bar: 20 μm); E, Metathorax setae 2T (scale bar: 10μm). Setae numbered according to the chaetotaxy proposed in study: 1P, 1T, 1M and 3T: forked-apex short trichoid sensilla (thick arrow); 2P and 2M: blunt short trichoid sensillum (thin arrow); * campaniform sensilla.

**Fig 13.**
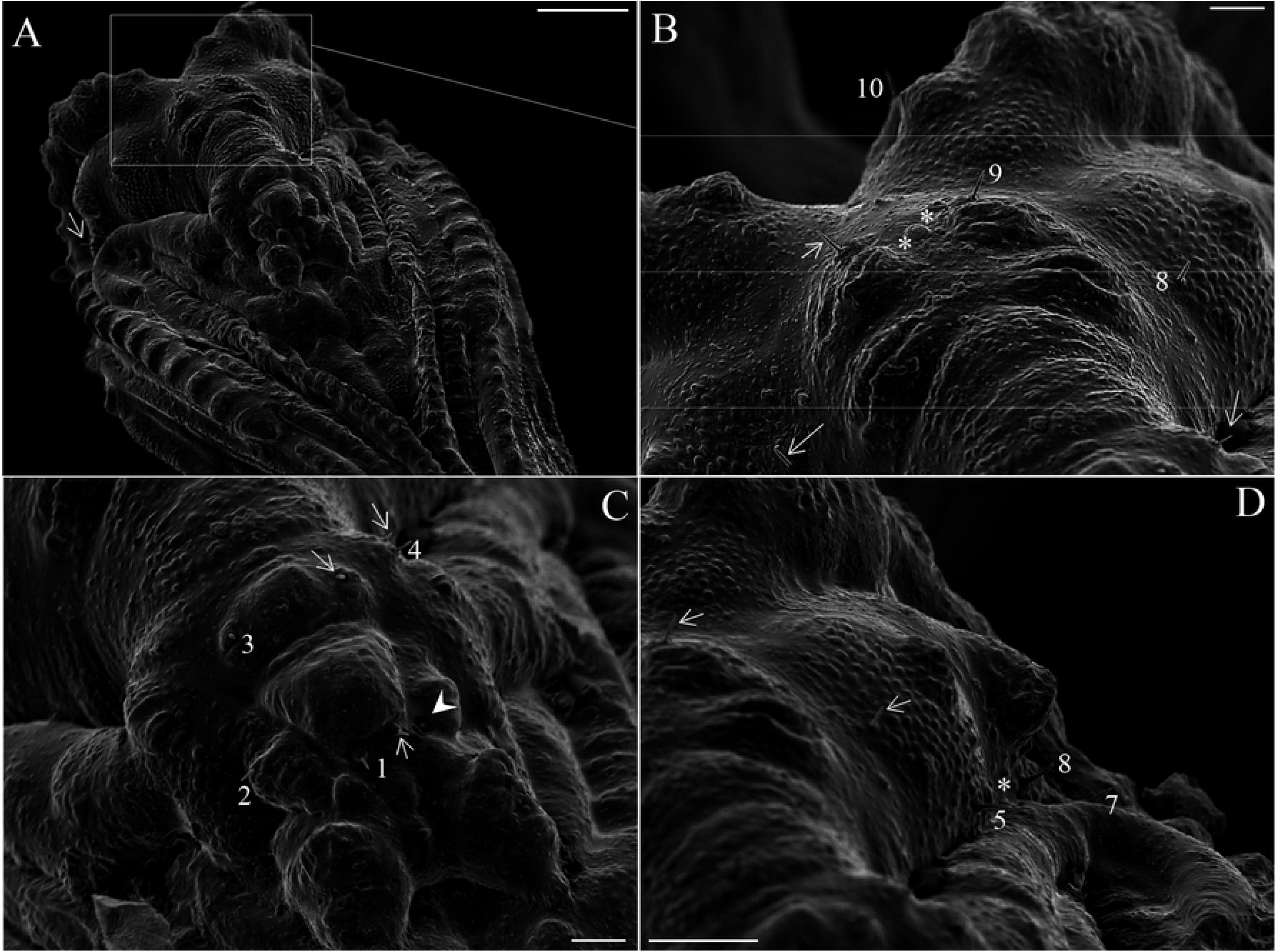
Scanning electron microscopy of the pupa of *Migonemyia migonei*. A, head of the pupa(scale bar: 100 μm).. -(B-D scale bars: 20 μm). Setae numbered according to the chaetotaxy proposed in this study. Clypeal inferior (1), Palpal seta (2), Superior clypeal (3), Inferior frontal (4), Medial postocular (5), Internal postocular (6), External postocular (7), Medial frontal (8), Superior frontal (9), Vertical (10).

The cephalic sheath with antennal impressions shows outlines of all flagellomeres of the pre-imaginal stage. Mouth part sheath is smooth; clypeal sheath is conspicuous; sexual dimorphism presented in the maxillary sheath is shorter than the sheath of the labrum-epipharynx and hypopharynx in the males; head chaetotaxy with very small spines (Fig. 13) and are numbered (Table 3). Thorax with a large longitudinal crest in the middle of dorsal side, Y-shaped, the pro and mesothorax have a pair of prominent tubercles, the methatorax has two pairs, the latter is bigger than the former (Fig. 12). There are also a pair of spiracles (ventilatory orifice per Oca-Aguilar et al., 2014) in the prothorax. The prothorax has 2 + 2 small setae in each one and a campaniform sensillae; mesothorax with 3 + 3 pairs of setae, one pair of them being long, chaotic, called prealar, not sharp at the tip, (length 0.15 ± 0.008 mm, n = 5) and are stout, originating from tubercles, and a large mesonotal tubercle with a continuous border (Fig. 12B); the mesotonal tubercle is considered here as part of the Y arm of the longitudinal crest, with scattered small rounded tubercles. Metathorax has four pairs of setae, some associated with tubercles, with a small bifurcation in the tip of each seta – 1T and 3T (Fig. 12 D and F). Ventral side of the thorax has leg and wing sheaths, and a marked wing venation stamped into it, with row of rounded tubercles in each stamped venation (Figs. 12A, 15A and 16A).

The tegument of the abdomen is covered by several small, spiniform tubercles. There are nine segments, and the width of every segment is near twice its own length. They diminish gradually in size towards the distal region (Fig. 10 A), becoming discrete lateral projections in the pleural sheath. The last segments show sexual dimorphism, with a posterior spiracle in the eighth segment. Segments I -VII have 2 pairs of median dorsal tubercles. Abdominal segments I-II have atergum with four pairs of setae, and pleura and sternum are covered with the thoracic appendage sheaths. Abdominal segment III has a tergum and sterna with 4 pairs of setae, respectively, on each side, which are similar in shape and location to the abdominal segments IV-VII; pleura and sterna III are partially covered by the thoracic appendage sheaths (Table 2). In abdominal segments IV-VII, each tergum has 4 pairs of setae which are distributed in a similar fashion to the previous segments. Each sternum has 4 pairs of setae. In abdominal segment VIII, males and females both have two pairs of setae on the tergo, and two pairs on the sternum and two pairs on the spiracle (Fig. 16F). All these are very small basiconic setae. Abdominal segment IX is covered by the larval exuvia (as is VIII), but when uncovered, sexual morphological differences can be observed. In males, there are two lobes on each side – one simple, covering the lateral lobe, and the other divided, containing the gonostylus and gonocoxite In females, two simple and short lobes on each side – one covering the oviscape and the other the cercus, though without setae (Fig. 17), and the genital opening sheath is discreet (Fig. 16E). Abdominal chaetotaxy can be seen in Table 2, Figs. 15 and 16.

**Fig 14.**
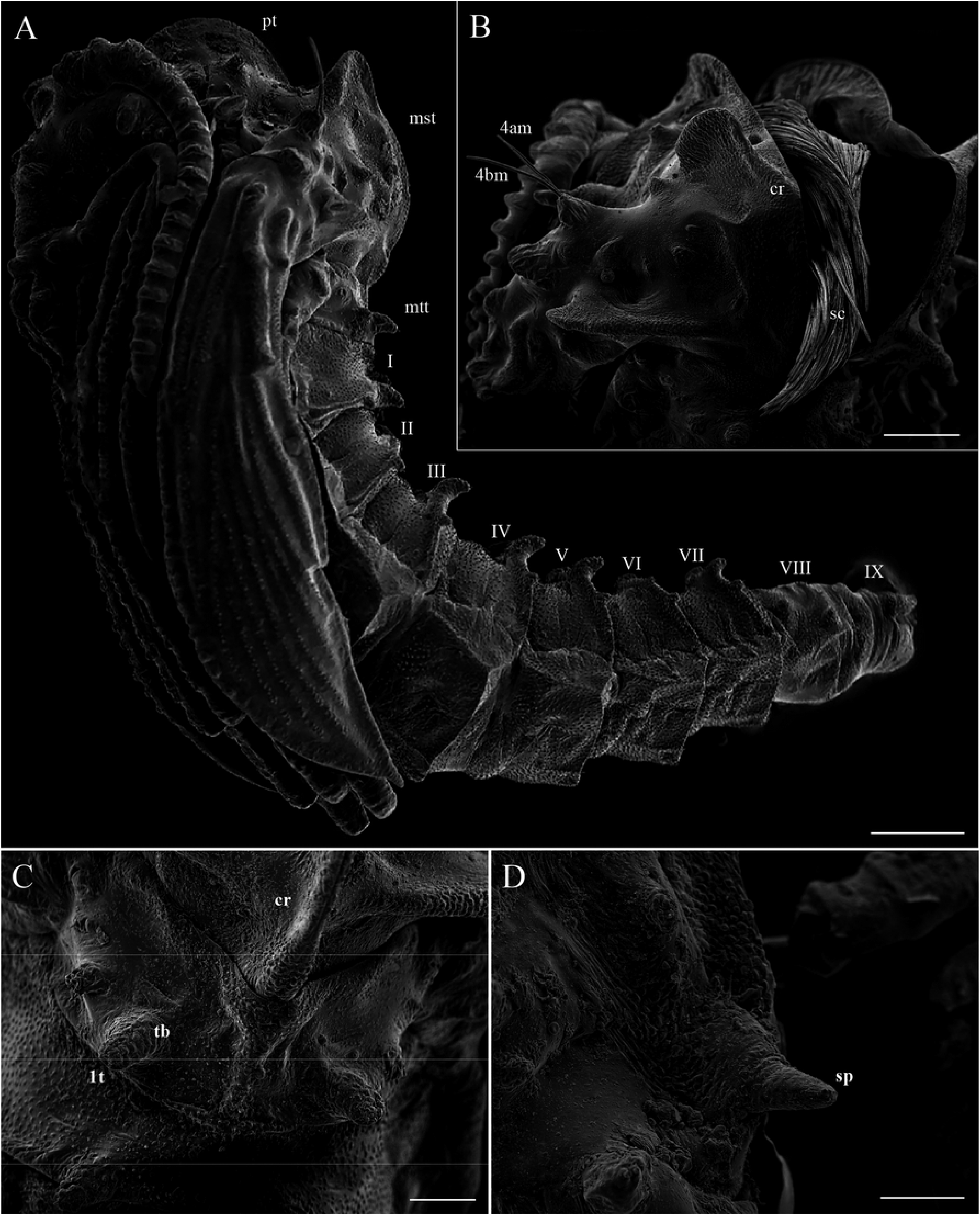
Scanning electron microscopy of the pupa of *Migonemyia migonei*. A, Pupa in lateral view. (scale bars: 200 μm); B, Opening of pupa crest, scale from the (scale bars: 100 μm); C, Metatorax the pupa; D, Superior spiracle pupal (C and D scale bars: 50 μm). Abbreviations: cr: crest, mst: mesotorax, mtt: metatorax, pt: protorax, sc: scales (hairs) of the pharate adult, sp: spiracle, tb: tubercle.

**Fig 15.**
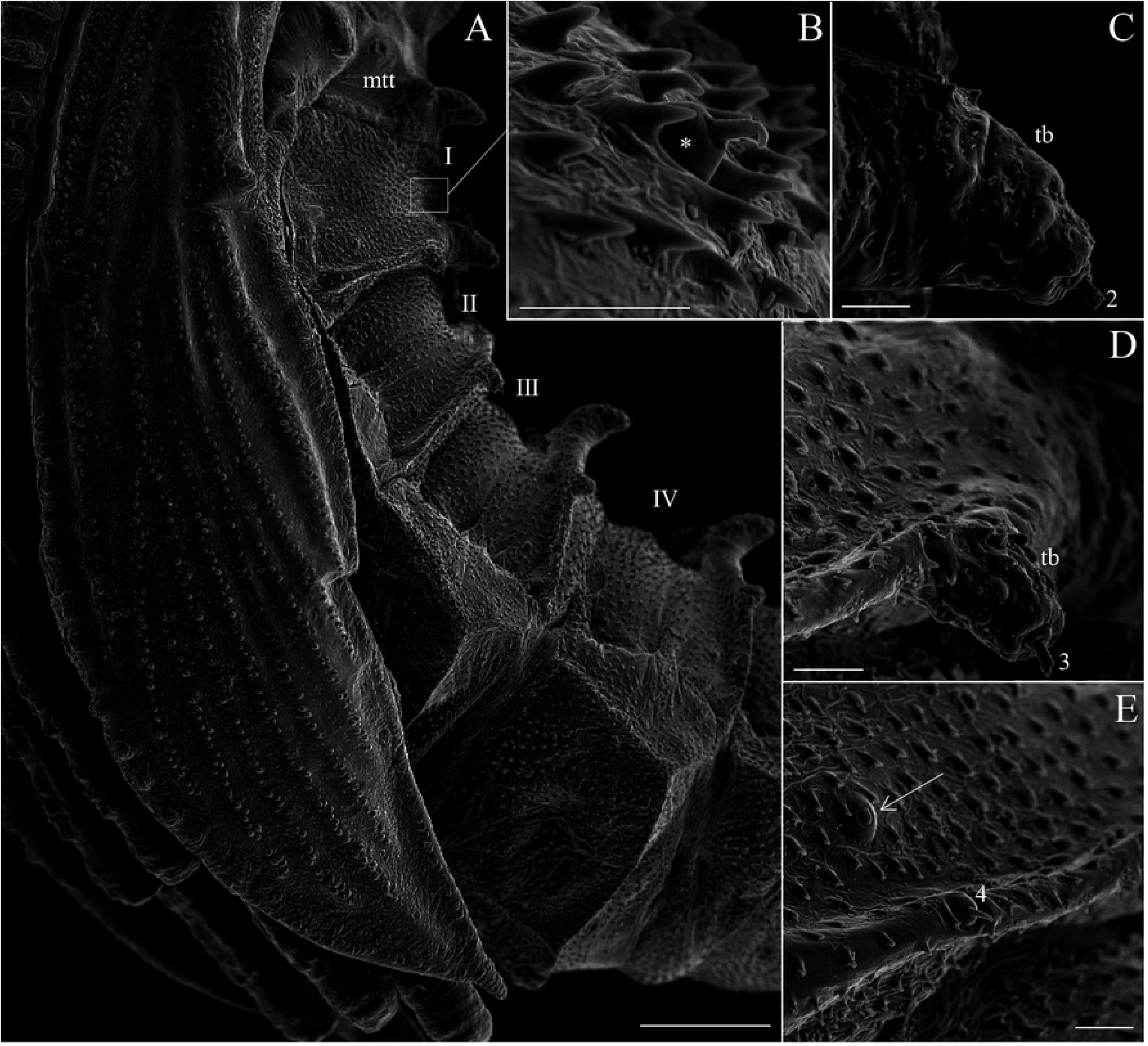
Scanning electron micrograph of the pupa lateral view of *Migonemyia migonei*. (scale bars of A: 100 μm, of B-D: 10 μm). As – abdominal segment, swv – sheath of wing venation, tb – tubercle, black arrow – campaniform sensilla, *increased area showing seta (1), black arrow pointing campaniform sensilla, Internal posterior dorsal (2), External posterior dorsal (3), Laterodorsal (4).

**Fig 16.**
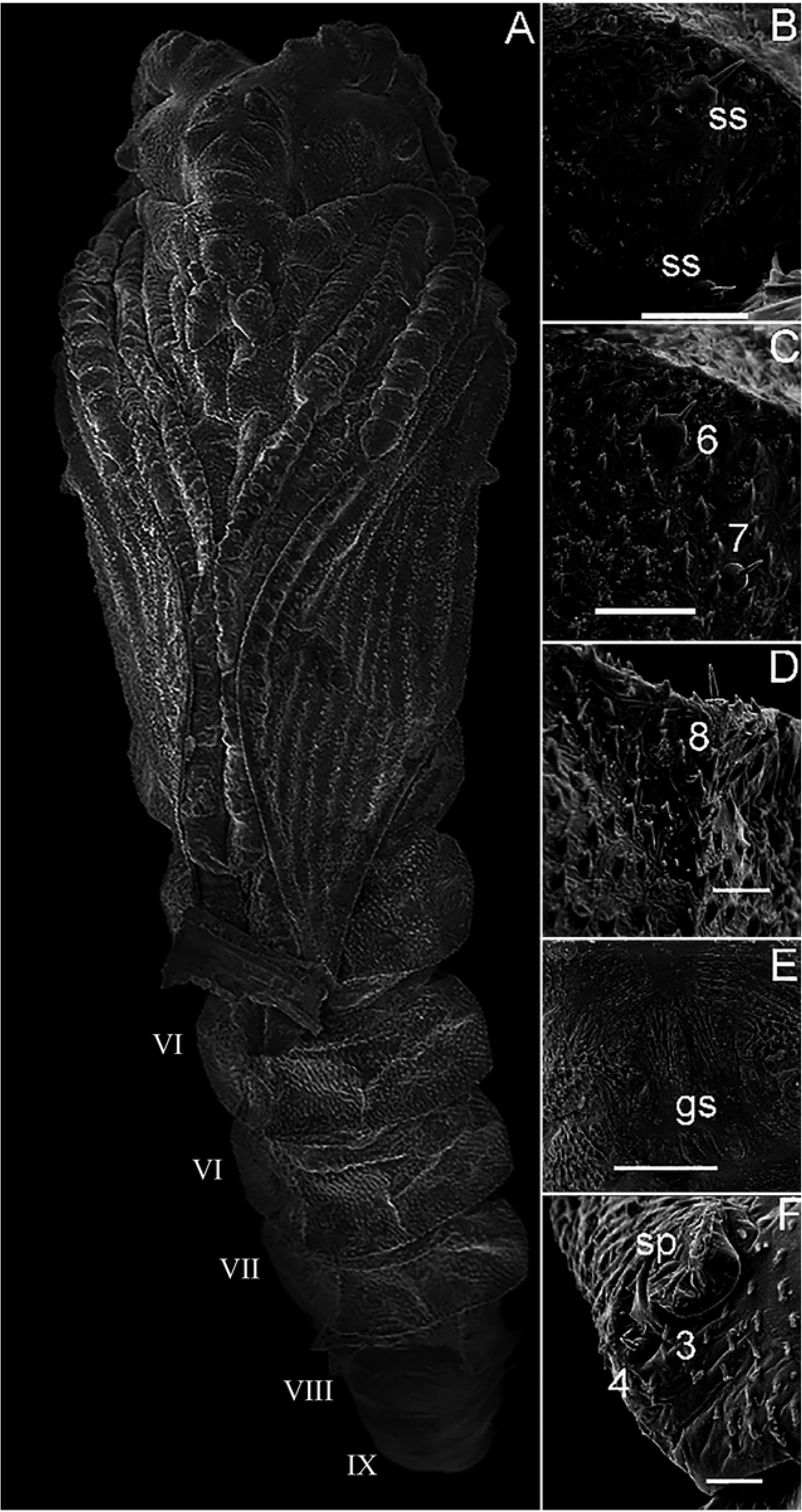
Scanning electron microscopy of the pupa of *Migonemyia migonei*. A, pupa ventral view; B, C and D ventral abdominal segment of setae; E, genital opening sheath; F, part of abdominal segment 8 with posterior spiracle. As – abdominal segment, gs – genital sheath, sp – spiracle, ss – External posterior dorsal (3), Laterodorsal (4), External posterior ventral (6), External anterior ventral (7), Internal posterior ventral (8).

**Fig. 17.**
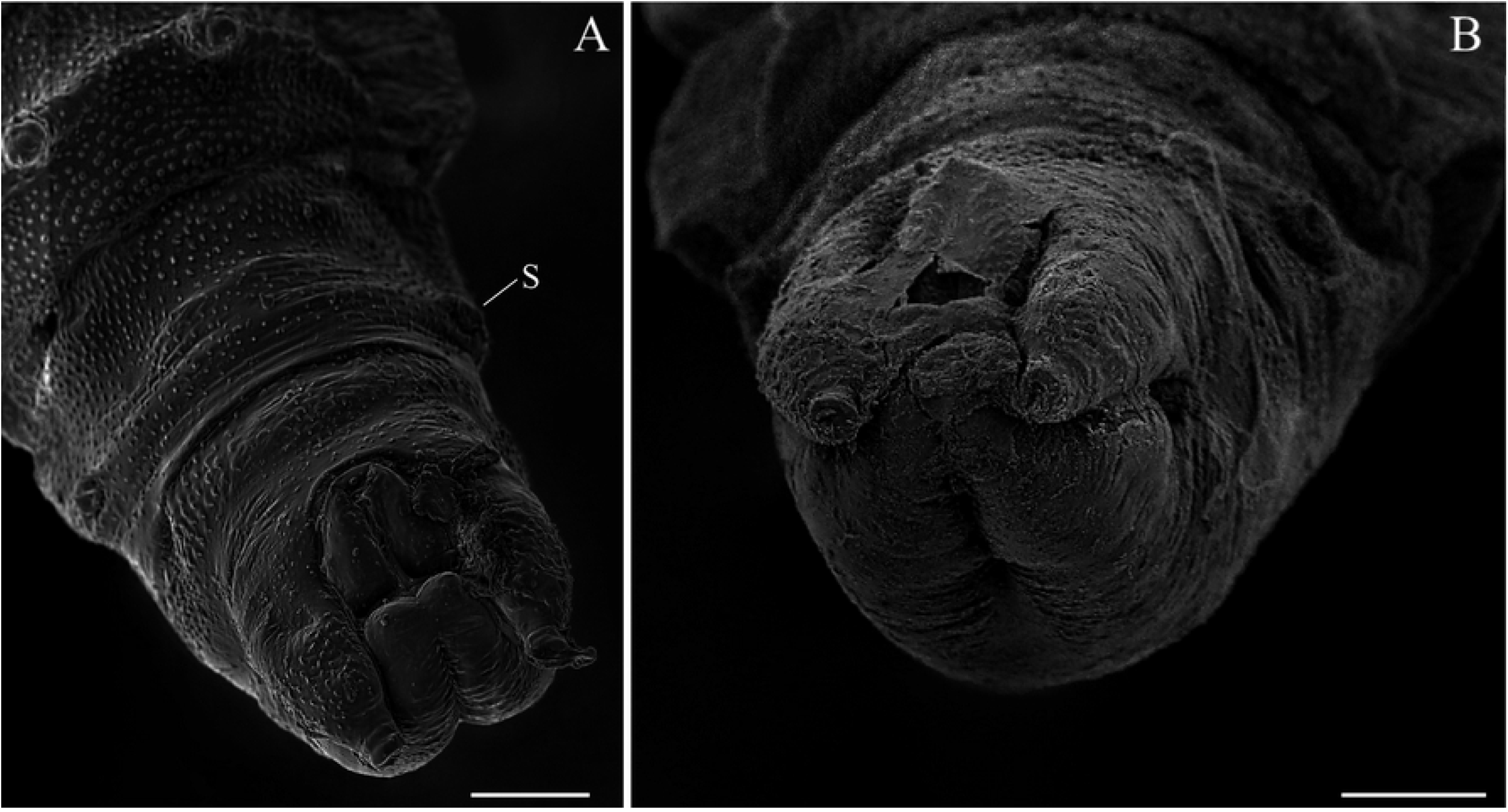
Scanning electron microscopy of the pupa of *Migonimyia migonei*. A, Pupa female in ventral view; B, Pupa male in ventral view (scale bars of A and B: 50 μm). s, spiracle.

**Table 2.** Chaetotaxy for pupa of *Migonemyia migonei*.

| Tagma | Number | Sensillum type | Terminology |
| --- | --- | --- | --- |
| HEAD | 1C | Basiconic | Clypeal inferior |
|  | 2C | Basiconic | Palpal seta |
|  | 3C | Basiconic | Clypeal superior |
|  | 4C | Basiconic | Frontal inferior |
|  | 9C | Basiconic | Frontal superior |
|  | 10C | Basiconic | Vertical |
|  | 8C | Basiconic | Frontal medial |
|  | 5C | Basiconic | Postocular medial |
|  | 6C | Basiconic | Postocular internal |
|  | 7C | Basiconic | Postocular external |
| THORAX |  |  |  |
| Prothorax | 1P | Basiconic | Protoracic superior |
|  | 2P | Basiconic | Prothoracic medial |
|  | 3P | Absent | Absent |
| Mesothorax | 1M | Styloconic | Mesothoracic inferior |
|  | 2M | Styloconic | Mesothoracic medial |
|  | 3M | Absent | Absent |
|  | 4A-B M | Chaetic | Pre-alar |
| Metathorax | 1T | Styloconic | Metathoracic internal |
|  | 2T | Styloconic | Metathoracic medial |
|  | 3T | Basiconic | Metathoracic external |
|  | 4A - B T | Basiconic | Pre-halter |
| ABDOMEN |  |  |  |
| I-VII | 1 | Basiconic | Dorsal anterior |
|  | 2 | Styloconic | Dorsal posterior internal |
|  | 3 | Styloconic | Dorsal posterior external |
|  | 4 | Basiconic | Laterodorsal |
|  | 8 | Basiconic | Ventral posterior internal |
|  | 7 | Basiconic | Ventral anterior external |
|  | 6 | Basiconic | Ventral posterior external |
|  | 9 | Basiconic | Ventral anterior internal |
| VIII | 1 | Basiconic | Dorsal superior |
|  | 2 | Basiconic | Dorsal inferior |
|  | 3 | Basiconic | Lateral |
|  | 4 | Basiconic | Lateral |
|  | 5 | Basiconic |  |
|  | 6 | Basiconic |  |

## Discussion

The exochorion pattern of *Mg. migonei* is polygonal, and was first described by Barreto[29], using only light microscopy and subsequently redescribed by Fausto[23] who used SEM. This polygonal pattern of exochorion sculptures is found in 28 other species of Neotropical sandflies [14,30], which are distributed in nine genera of different subtribes. This characteristic probably has a phylogenetical importance, according to the opinion of Perez & Ogosuku [31], who state that the exochoronic pattern does not reflect phylogenetic relationships based on adult characteristics. Ward and Ready [32] and Costa [33] suggested that the design of the sand fly’s exochorion could be different according to the environment of the breeding site. Bahia [7] observed through SEM that the eggs of *Ny. intermedia* and *Ny. whitmani* presented a different exochorion pattern than that observed in *Mg. migonei*. Instead of presenting ornaments with polygonal reticulation, with alternating transversal rows of generally rectangular parallel cells or square to polygonal cells, the eggs of these species of *Nyssomyia* presented ornamentations that consist of parallel ridges covering the entireexchorion. This exochorion pattern was previously observed in these same species through light microscopy by Barretto [29], who described it as “connected ridges”. Subsequently, Pessoa [12] described the chorion of *Ev. Carmelinoi*, and the genus *Evandromyia* is phylogenetically close to *Migonemyia*. Nevertheless, the knowledge about sand fly zootaxonomy is a little scarce.

The chaetotaxy and morphological structures of Neotropical phlebotomine larvae have been discussed previously by several authors [4,7,8,12,14–16,27,29,32,34–38]. The general aspect of head, the position of the mouthparts a somewhat prognathous, the peculiar shape of the egg buster, similar to a “volcanic cone” [7], the body shape cylindric, and the types of the most part of the bristles observed in *Mg. migonei* larvae have a similar pattern to that observed by SEM in larvae of the sand fly species *Nyssomyia intermedia* and *Ny. whitmani* [7] and also of some larvae from species grouped in the Lutzomyiina and Sergentomyiina subtribes e.g. *Lutzomyia longipalpis, Lu. cruciata, Ev. carmelinoi* [12], *Micropygomyia chiapanensis* larvae description [4].

Trichoid sensilla, of different subtypes, are the most common types found on larvae and adults of phebotomine sand flies, among others Diptera [7,39,40]. The short and long trichoid sensilla evidenced in the present work surrounding the base of the anal lobes and on the lateral sites of the prolegs of *Mg. migonei* larvae are similar to those previously observed in the same site of *Ny. intermedia* and *Ny. whitmani* larvae [7]. Other similarities in relation to the typology and the pattern of sensillary distribution were also evidenced among the larvae of these species, such as brush-like trichoid sensilla, also located in front of the egg burster and on the lateral and dorsal aspects of the body segments, trichoid sensilla on the apex of the head, short trichoid sensilla on the mouthparts and weakly brush-like trichoid sensilla inserted slightly forward and between the antennae, and long trichoid sensilla are inserted further down towards the mouthparts.

In addition to these sensillary types, other similar sensilla such as one apical clavate basiconic sensillum, on the apex of the antennae of the *Mg. migonei* larvae, and one clavate coeloconic sensilla and three short blunt coeloconic sensilla, implanted on the proximal region of antennae, were also evident in *Ny. intermedia*, and *Ny. whitmani* larvae [7]. The latter sensillary subtype, when observed in higher magnification by Bahia [7] in larvae of *Ny. intermedia*, showed to have wall pores (multiporous clavate coeloconic sensilla; a SW-sensillum subtype). A similar sensory subtype, presenting wall pores, was also identified at the equivalent larvae site of *Lu. longipalpis*, by Pessoa [11], previously designated as “multiporous papilla”. Sensilla similar to the apical clavate basiconic sensillum and the clavate coeloconic sensilla were also evidenced in the larvae antennae of the *Lu. cruciata* sand fly by Oca-Aguilar [15].

The antennal pattern of mature larvae of *Mg. migonei* seen under SEM was done by Pessoa [11], and the other earlier stages described here can be included in category iv of Leite and Williams [5] proposal of antennae shape. To mouthpart, it is possible to highlight the teeth of mandibles because of the position of the mandible, ordinarily described in the lateral view. The chaetotaxy the thorax of *Mg. migonei* is quite similar to *Ev. carmelinoi* and *Ev. lenti*, a genus close to *Mygonemyia*, and which was described using similar methodology. In the thorax, only the shoulder accessory b seta is evidently different, very small, bifid or trifid and, in *Mg. migonei*, it is a double the size, semi-barbed, while absent in *Ev. lenti*.

The anterior spiracles of the *Mg. migonei* population obtained from the Ceará State in Brazil possess a few more papillae (8-9) than those from the Mérida state in Venezuela (7)[9]. *Ev. carmelinoi* and *Ev. lenti* also have 8[12]. The number of papillae of the posterior spiracle are similar between the two populations of *Mg. migonei* and *Ev. carmelinoi* and *Ev. lenti*. The dorsal submedian setae 7 and 8 of the meso and metathorax and abdominal segments I-VII of *Mg. migonei* are similar in size to *Ev. Carmelinoi*[12], and both have the double the size of setae compared to *Ev. lenti*[12]. The setae 11 and 12 of *Mg. migonei* are barbed and have the double the size of the same setae in *Ev. carmelinoi* and *Ev. lenti[12]*, which in their case are simple. The last segment also has slight differences; t setae 11 in *Mg. migonei* are bigger than *Ev. Lenti*, and seta 12 in *Mg. migonei* is only half the size of its counterpart in *Ev. lenti* [12]. A large campaniform sensillae present in the ventral side of each tubercle of implantation of the caudal setae had not been described before and it is duly registered here.

The caudal filaments evidenced in the last larval segment of *Mg. migonei* and in other sand flies, is a long subtype of trichoid sensilla, which presents multiporous wall (SW-sensilla), being classified, therefore, as an olfactory sensory structure [41,42]. Olfactory sensilla presents in their surperficial microstructure particular very noticeable characters: multiporous walls (SW-sensilla) or walls with longitudinal grooves (DW-sensilla) [42–44]. However, these characteristics determine the generic olfactory function. The response to which odor molecules respond to each sensilla can only be determined using electrophysiological bioassays, especially the Single Sensillum Recording coupled to Gas Chromatography (SSR-GC),Single Sensillum Recording coupled to Gas Chromatography (SSR-GC), an efficient method for isolating potential insect attractants, the action potentials of odor receptor neurons (ORNs) present in each type of olfactory sensilla can be recorded *in situ* [45]. In this sense, we highlight the great importance of carrying out previous studies by scanning electron microscopy to identify the olfactory sensilla and indicate the sensillary topography, the precise location of these sensory structures, to support the performance of bioassays with SSR-GC, facilitating the accurate orientation and targeting of the electrodes and the odours pulses tested, especially in analysis of insects with very small antennas and very covered by pilosities, such as sand flies among other Diptera, e.g. Different species of sandflies have different pore patterns in their caudal filaments. *N. whitmani* larvae, for example, presents in their caudal filaments pores distributed within wall grooves, deep and not very close, whereas *Ny. intermedia’s* have more separate and superficial pores, not found in well-defined grooves [7]. *Ev. lenti* larvae have a smaller number of pores in their caudal filaments, distributed along longitudinal, thin, parallel and closer ridges [11]. The caudal filaments of *Mg. migonei* larvae, in turn, presents a larger number of pores, deep, between non-parallel ridges, but that interconnect. We believe that the differences observed between the distribution patterns of the pores of the caudal filament wall may serve as important characters in future taxonomic and phylogenetic studies with larvae, for a better understanding of the evolution, of the allopatric speciation process in Phlebotominae.

We presented in this study the first images of the pupa emerging from the Y suture of the head, and subsequently, the pupa, by movements of contraction and inflation, ruptures the thorax in the middle of the dorsal part. There are only a few pupae from Neotropical sand fly species described in detail using 3D images of SEM, howver, all the head setae, small spines, and basiconics are homologous to other Neotropical pupae described [8,13–16]. Nevertheless, some minor differences can be highlighted in the thorax. The setae 1P and 2P are basiconic, 1P is bifid and stout and implanted in a tubercle. In some species, the setae 1P and 2P are styloconic, or at least one of them is, e.g*., Da. beltrani[8], Mg. chiapanensi[14], Lu. cruciata[15], Ny. umbratilis[16]*, and this characteristic is not determinant to phylogeny for Lutzomyiina. The absence of the 3P seta in *Mg. migonei* is probably an apomorpy in this species, at least in comparison to the others discussed here in this paper. The abdominal segments I -VII have 2 pairs of median dorsal tubercles, one large and conspicuous curved backwards. *My. chiapanensis[14]* and *Lu. cruciata[15]* possess these tubercles, however they are discrete, and in *Da. beltrani*[8] and *Ny. umbratilis*[16] apparently they are not in evidence. The setae 5, absent, is present in most pupae described, except for *Ny. umbratilis* [16].

In the present study, the immature stages of *Mg. migonei* possess some discrete or evident structures that can be used as apomorphies for phylogenetical relationships in order to understand the evolutionary history of this species. *Migonemyia migonei* is possibly a species complex [46]. It is expected that the present descriptions may contribute to the taxonomy status, at least for that of *Mg. migonei* from the Ceará population, which occurs in an important endemic area for cutaneous leishmaniasis in Brazil.

